# Exposure to novel females increases fecundity in adult male prairie voles

**DOI:** 10.1101/2024.12.18.627569

**Authors:** Jessica A. Hurd, Yurika L. Watanabe, Gracie J. Toben, Alexandra K. Ford, Craig A. Miller, Casey E. Sergott, Dale Kelley, Zoe R. Donaldson, Elizabeth A. McCullagh

**Author notes:** **Corresponding Author:** Jessica A. Hurd –. Yurika L. Watanabe –, Gracie J. Toben –, Alexandra K. Ford –, Craig A. Miller –, Casey E. Sergott –, Dale Kelley –, Zoe R. Donaldson –, Elizabeth A. McCullagh –.

## Abstract

Social circuitry of the mammalian brain can influence male reproductive physiology. This often manifests as plasticity in sperm production or allocation, particularly in response to male-male competition. However, socially mediated testicular plasticity has not been investigated with respect to mating and parental strategy. Testis mass and sperm production of sexually naïve and female-exposed adult male individuals of three rodent species were compared: the socially monogamous and paternal prairie vole (*Microtus ochrogaster*), the promiscuous and non-paternal meadow vole (*Microtus pennsylvanicus*), and the promiscuous and non-paternal house mouse (*Mus musculus*). Monogamously paired prairie vole males exhibited significantly larger testes and greater sperm production than naïve prairie vole males. Comparatively, there were no significant differences between naïve and monogamously paired male meadow voles or mice. To investigate the role of olfactory cues for regulating this phenomenon in prairie voles, a group of naïve males exposed to soiled bedding from novel females was used. These males were more similar to paired males than to naïve males not exposed to novel female odors, demonstrating a strong role of the social olfactory system. Further, while the predictions of sperm competition theory (species with greater female promiscuity have larger testes than closely related species with less female promiscuity) are consistent between naïve meadow voles and prairie voles, the prediction does not hold for monogamously paired prairie voles and meadow voles. This demonstrates the complexity of internal social dynamics and reproductive pressures which socially monogamous paternal males face and the evolutionary adaptions that may develop in response.

## Introduction

The social circuitry of the mammalian brain can impact an animal’s physiology, behavior, neurochemistry, and neuroanatomy from both evolutionary and life-history perspectives (Blumenthal & Young, 2023; Ferreira-Fernandes & Peça, 2022; Lukas & Clutton-Brock, 2013; Nelson, 2015; van der Horst & Maree, 2014). In the mammalian reproductive system, primary mating strategies and prevalence of sexual promiscuity in a species often correlates to trends in reproductive physiology (Kramer & Russell, 2015; Lukas & Clutton-Brock, 2013; Ophir & delBarco-Trillo, 2007; Schacht & Kramer, 2019; van der Horst & Maree, 2014). For example, there is an evolutionary pattern among male mammals that differences in relative testis size between species serve as an indicator of general female promiscuity in the species, with low rates of sperm competition resulting in smaller testes relative to body size (Kramer & Russell, 2015; Schacht & Kramer, 2019; van der Horst & Maree, 2014). This interspecies, evolutionary perspective, however, fails to address how the social environment and social structure may cause differences within and between male individuals of the same species from an intraspecific life history perspective. Within the individuals of a species, social encounters and the extrinsic social environment can impact gonad function, partially via the hypothalamic-gonadal-pituitary (HPG) axis (C. Carter, 1987; C. S. Carter et al., 1980; Dixson & Anderson, 2004; Firman et al., 2018; Koyama & Kamimura, 2000; Lukas & Clutton-Brock, 2013; Maruska & Fernald, 2011a; Ramm & Stockley, 2009a; Taylor et al., 1986). Further, in many mammal species, the social environment can impact testicular function and/or sperm quality (Dixson & Anderson, 2004; Firman et al., 2018; Koyama & Kamimura, 2000; Lukas & Clutton-Brock, 2013; Maruska & Fernald, 2011a; Ramm & Stockley, 2009a). Social triggers of these changes often involve male-male competition (Maruska & Fernald, 2011a; Ramm & Stockley, 2009a), but may also include things such as sexual experience (Taylor et al., 1986) and parental status (Bales & Saltzman, 2016; Reburn & Wynne-Edwards, 1999). However, how the social environment alters testicular function on a molecular level is poorly understood (Maruska & Fernald, 2011a).

Most work investigating the impacts of the social environment on male reproductive function in mammals has been done in promiscuous, non-paternal species. Little investigation has been done on this topic in socially monogamous mammals, which make up ∼5-9% of the class (Blumenthal & Young, 2023; Lukas & Clutton-Brock, 2013; van der Horst & Maree, 2014). While paternal care is rare in mammals, it occurs in 56-59% of socially monogamous species (Lukas & Clutton-Brock, 2013; Stockley & Hobson, 2016). Social monogamy is sometimes erroneously correlated with sexual fidelity, which does not reflect the true nature of many socially monogamous species which frequently exhibit variable rates of extra-pair sexual encounters (Lambert et al., 2018; Ophir et al., 2008; Young, 2003). The presence of both a socially monogamous, biparental mating style as well as variable sperm competition creates an intriguing situation regarding male reproductive function and plasticity. Social monogamy provides reliable access to a consistent sexual partner (Lambert et al., 2018) and paternal care can increase reproductive success (Stockley & Hobson, 2016). On the other hand, parenting offspring that may not be one’s own due to extra-pair sexual encounters is costly from a fitness perspective. This generates a need for paired males to reconcile the convenience of a guaranteed sexual partner while also combating paternity loss.

Prairie voles (*Microtus ochrogaster*) – a hamster-sized, burrowing rodent native to North American prairies – may serve as an excellent model species for investigating the impacts of the social environment on male reproductive function in socially monogamous, pair-bonding, biparental mammals (Young, 2003). In addition to being socially monogamous, prairie vole males are highly paternal (Getz et al., 1981; McGraw & Young, 2010) and even alloparental (Finton et al., 2022; Roberts et al., 1998). Female prairie voles exhibit social modulation of their reproductive organs and behavior. They are induced ovulators and will remain in an anestrus state until exposed to an unfamiliar conspecific male (C. Carter, 1987). They also exhibit a direct familial barrier to reproduction and will not normally ovulate when exposed only to their fathers and male littermates (C. S. Carter et al., 1980). On the contrary, little is known about how the social environment affects testicular function in male prairie voles and how those effects may be specifically related to the prevalence of social monogamy and paternal care exhibited by the species. Prairie voles offer the opportunity to study the relationship between mating and parental strategies and reproductive physiology in a controlled, laboratory setting.

To elucidate the relationship between female exposure and testicular plasticity within a limited monogamy and promiscuity framework, we will compare the effects of novel female exposure on testicular function in three rodent species – the socially monogamous and biparental North American prairie vole, the closely related promiscuous and nonpaternal North American meadow vole (*Microtus pennsylvanicus*), and the unrelated promiscuous and nonpaternal house mouse (*Mus musculus*). Sperm competition can alter reproductive function in rodents, including house mice and meadow voles (delBarco-Trillo & Ferkin, 2006; Firman et al., 2018; Ramm & Stockley, 2009a). To our knowledge, the effects of sperm competition on prairie vole reproductive function have never been investigated. Because house mice and meadow voles do not form pair bonds nor provide paternal care, we hypothesize that female pairing alone in the absence of male competition will not induce any changes in testis size, sperm counts, or testicle histology in these species. In contrast, we hypothesize that male prairie voles will exhibit an increase in testis size and sperm production solely in response to female exposure due to the reproductive pressures generated by social monogamy and paternal care.

## Materials and Methods

To investigate the effect of female pairing on testis mass, testis composition, and sperm production, same-sex housed, sexually naïve males were compared to monogamously housed, female-paired males in all three species: prairie voles (*Microtus ochrogaster*), meadow voles (*Microtus pennsylvanicus*), and house mice (*Mus musculus*). One additional experimental group of prairie voles was also used, consisting of naïve males exposed only to soiled female bedding, dubbed the olfactory exposure (OE) group. Fecal testosterone metabolite data was collected on all three prairie vole groups.

### Ethical Note

All experiments complied with all applicable laws, National Institutes of Health guidelines, USDA guidelines, and were approved by the Oklahoma State University IACUC, approval number 22-09. Investigators understand the ethical principles under which the journal operates and that their work complies with the animal ethics checklist.

### Subjects

#### Prairie Voles

Prairie voles were laboratory reared and maintained at Oklahoma State University, originally obtained in 2020 from Dr. Tom Curtis’s colony at the Oklahoma State Health Sciences Campus. The light:dark cycle was 14:10. Animals were weaned at ∼postnatal day (PND) 21 and housed with 1-2 same-sex animals until harvest or pairing. All animals were between 90-300 days old at the time of harvest.

#### Meadow Voles

Meadow voles were laboratory reared and maintained at University of Colorado Boulder, originally obtained from research colonies at Smith College and University of California San Francisco. The light:dark cycle was 14:10. Animals were weaned at ∼PND21 and housed with 2-4 same-sex animals until harvest or pairing. All animals were between 100-720 days old at the time of harvest.

#### House Mice

C57BL/6J (stock #000664, B6) wildtype background mice were laboratory reared and maintained at Oklahoma State University, originally obtained from the Jackson Laboratory (Bar harbor, ME USA). The light:dark cycle was 12:12. Animals were weaned at ∼PND21-30 and housed with 1-4 same-sex siblings until harvest or pairing. All animals were between 90-450 days old at the time of harvest. In addition to examination as an evolutionary outgroup, house mice are useful to help evaluate the validity of the utilized sampling methods.

### Experimental Housing for all Animals

Animals were housed in a climate and light controlled housing environment with consistent parameters year-round. Food and water were provided ad-libitum, along with regularly changed bedding and nesting material, and enrichment as needed. All animals were harvested at a minimum of PND95.

### Procedures

#### Female Pairing

For the paired males, pairing occurred at a minimum of PND60 for all species. Varying lengths of pairing were included, with the shortest being long enough to ensure at least one full cycle of spermatogenesis. This is ∼29 days in prairie voles (Schuler & Gier, 1976) and ∼35 days in house mice (Mäkelä & Toppari, 2018; Oakberg, 1956). The length of the seminiferous epithelium and spermatogenesis cycles have not yet been characterized in meadow voles. Therefore, a minimum of 6 weeks of pairing were provided, as this is reasonably beyond the length of spermatogenesis reported in three other vole species – 29 days for prairie voles, 37 days for *Microtus agrestis*, and 31 days for *Clethrionomys glareolus* (Grocock & Clarke, 1976; Schuler & Gier, 1976). Social housing for each breeding pair consisted of the adult male and female and their pups with no additional adult animals present, ensuring a lack of direct sperm competition. Naïve males were not exposed to any direct or indirect encounters or cues from any females or males other than their own cage mates at any time.

#### Female-Soiled Bedding Exposure (OE prairie vole group only)

Experimental conditions were introduced at a minimum of PND70. 20-30g of soiled bedding from a cage of 2-3 unfamiliar, conspecific females was placed in the male cage twice weekly, 3-4 days apart for a minimum of 29 days to ensure at least one complete spermatogenic cycle before harvest. During this time, the cage of females also received 20-30g of soiled bedding from the cage of OE males to encourage an increase in uterine weight and potentially some resulting variation in fecal hormone profiles (C. Carter, 1987). Soiled bedding transfers were always performed between the same two cages of males and females so the males would not experience new perceived sperm competition from novel conspecific males via bedding transfer (delBarco-Trillo & Ferkin, 2006).

#### Testis Weight and Sperm Quantification

The protocol for testicle harvest and sperm cell removal was inspired by an artificial insemination protocol by Duselis and Vrana (Duselis & Vrana, 2007) and further modified for data analytic purposes. At the time of harvest, all males were euthanized with an lethal dose of isoflurane. The testes and epididymides were removed, separated, trimmed of fat, and weighed. The testes were transferred to 10% neutral buffered formalin (NBF) for preservation and storage. The epididymides were transferred to a small, conical-shaped container holding 0.5mL of 1x phosphate buffered saline (PBS) solution warmed to 37°C and left to acclimate for 5 minutes. They were then shredded with 18-gauge needles to sufficiently destroy the tissue and release the sperm cells into solution. The shredded epididymal tissue was allowed to rest in the PBS solution for an additional 5 minutes and then viewed via standard bright-field microscopy to visually confirm that the sperm cells diffused from the epididymal tissue. The solution was transferred to a plastic microtube containing 0.5mL of 10% NBF. This results in a final solution of sperm cells suspended in 1mL of 5% NBF, which effectively stores sperm cells for extended periods of time (Grønstøl et al., 2023; Schmoll et al., 2016). From this stored solution, total sperm counts were quantified with a NucleoCounter® (model SP-100) from Chemometec (Allerod, Denmark) at a dilution factor of 101 to reduce human counting error or bias.

For all prairie voles and house mice, the testes and sperm harvest began immediately after euthanasia. Since the meadow voles were housed with the collaborating laboratory at UC Boulder, the team removed, packaged, iced, and overnight-shipped the gonads from the experimental meadow voles to OSU for sperm extraction, histology, and data analysis. Cauda epididymides were immediately placed into a microtube with 0.5mL of PBS prior to shipping preparation. Upon reception, samples were unpackaged and warmed to room temperature. The testes and epididymides were trimmed of fat and processed according to the protocol described above. Instead of using 0.5mL of fresh PBS, the epididymides were shredded in the 0.5mL of PBS in which they were shipped, after being warmed to 37°C. This prevented shipping-related loss of epididymal sperm cells, which will remain suspended in the shipped solution.

#### Testicle Histology

Stored formalin-fixed testes were sent to the OCRID Immunopathology Core (IPC) at Oklahoma State University to undergo blind sectioning, staining, and scoring. Tissues were trimmed and processed in an automatic tissue processor (Leica ASP300s Tissue Processor, Leica Biosystems, Deer Park, IL USA). A delayed short cycle protocol was used, consisting of tissues held in 10% NBF, dehydrated in graded ethyl alcohols, cleared in toluene, and impregnated with paraffin (Leica Surgipath Paraplast Infiltration and Embedding Medium; Leica Biosystems). Tissues were embedded with the same paraffin into metal molds. Four to five µm sections were cut and collected onto charged slides (VWR #16004-408, Radnor, PA USA). Slides were dried overnight at room temperature and stained in an automated H&E stainer (Sakura Finetek DRS601, Torrance, CA USA). Microscopic evaluation of testes was performed by two board-certified veterinary pathologists. Categories that were scored included number of Leydig cells, Sertoli cells, seminiferous tubules, and premature germ cells. An ordinal scoring system was used where 0 = none, 1 = minimal, 2 = mild, 3 = moderate, and 4 = marked.

#### Fecal Testosterone Metabolites

Fecal testosterone metabolites were analyzed on different individuals than those used for testis and sperm data, as the decision to collect testosterone data was made after harvesting the initial set of animals. No sperm or testis data was collected on the males from which feces were gathered.

Fecal testosterone metabolites were quantified via the Arbor Assay (Ann Abor, MI USA) Detect X Testosterone H032-H1 Kit (Sensitivity of 15.2pg/mL). As this kit has never been previously utilized in prairie voles, fecal extractions were performed on preliminary samples using either 60% ethanol or 80% methanol (Palme et al., 2013), followed by a serial dilution experiment. Both alcohols provided similar results and demonstrated dilution linearity, therefore 60% ethanol was utilized for the experimental samples. The extracted samples were diluted in assay buffer at a dilution factor of 12.1 and the kit was used according to manufacturer instructions. Samples were run in triplicate with standards and controls in duplicate. For samples with an intraassay % coefficient of variation (CV) >10, the far-off triplicate was excluded and %CV re-calculated with the remaining two values. All 24 samples exhibited a resulting %CV of <16 and thus all were kept for analysis. All samples used for this analysis were extracted simultaneously and used together on the same plate in the same assay experiment. Thus, batch effect was avoided.

#### Statistical Analysis and Figure Generation

Data was analyzed in RStudio version 2024.9.0.375 (Posit Team, 2024) via linear regression (included in R Studio) and package emmeans (Length, 2024). Potential effects of age, weight, and length of female exposure on testis size and/or sperm count were examined via linear regression individually for each experimental group.

Differences in summed testis weights, sperm counts, histology variables, and testosterone data between experimental groups were then analyzed. Linear regression was performed for multi-species comparisons with pairing status and species as the predictors, in-line with a best-fit analysis determined beforehand via ANOVA. Estimated Marginal Means (package emmeans) – was used for posthoc pairwise comparisons between naïve and paired individuals within each species. Comparisons between the three experimental prairie vole groups were made via linear regression with female exposure status as the predictor. Posthoc pairwise comparisons between female exposure groups were performed via emmeans. Significance was determined by an alpha cutoff of 0.05. Means in text and tables are represented with ± standard error.

Graphs were also generated in R Studio with ggplot2 (Wickham, 2016). Significance bars were created with ggsignif (Ahlmann-Eltze & Patil, 2021), but significance values were annotated manually based on the generated statistical files described above. In figures 1, 2, 5, 6, 9, and 10, violin plots with individual data points demonstrate the full data distribution for each group. Means are represented by navy-colored horizontal bars. The black error bars represent standard error. In Figures 3 and 7, grouped bar graphs indicate the means. Errors bars represent standard error. For all figures, p-values of <0.05 are represented with “ * “, <0.01 with “ ** “, and <0.001 with “ *** “. P-values between 0.05 and 0.1 are not considered significant but may represent weak trends and are thus labeled on figures with the precise values. P-values >0.1 are labeled “N.S.”

**Figure 1:**
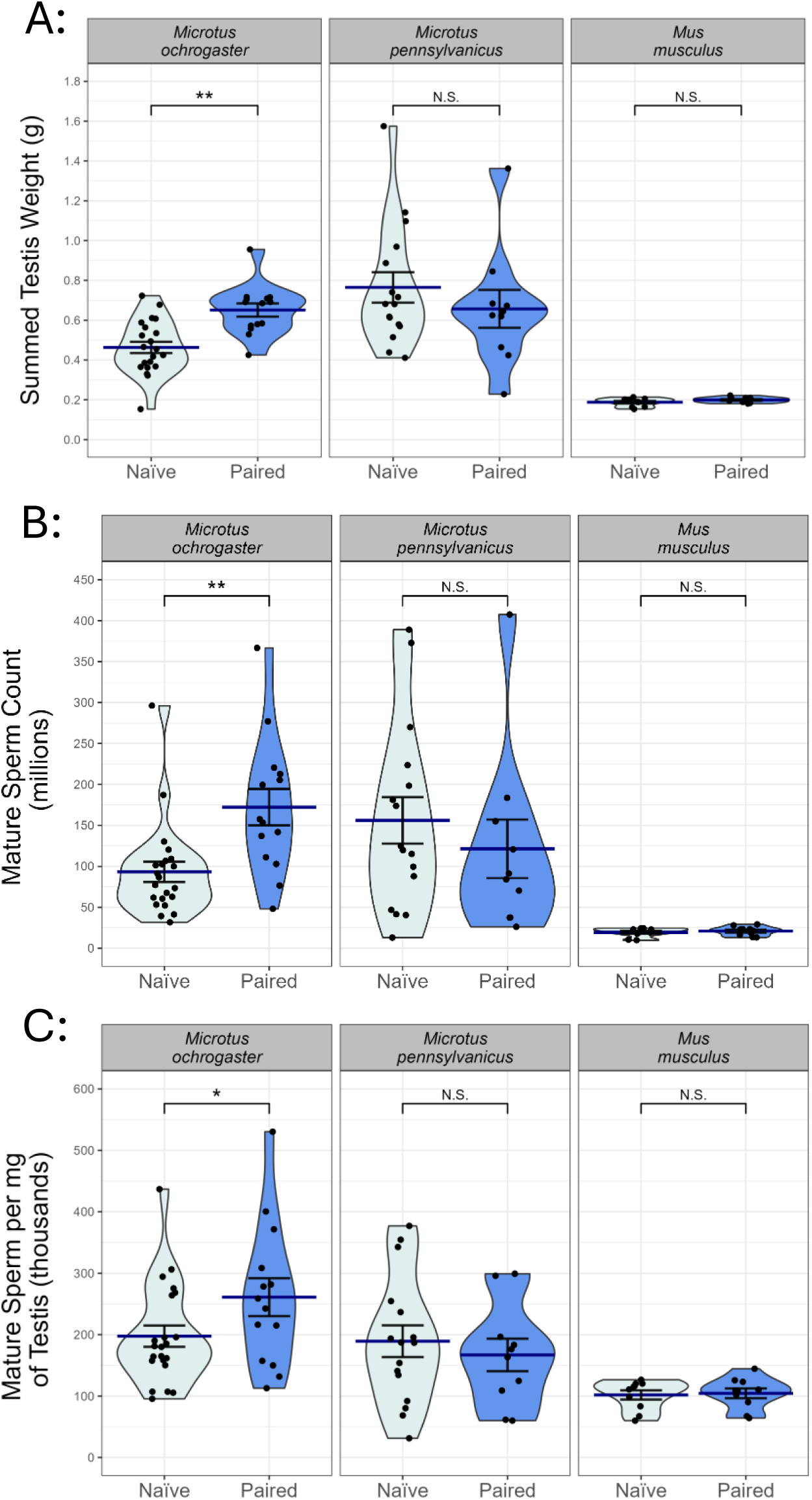
Comparison of A) summed testis weight (g), B) mature sperm count (millions), and C) mature sperm per mg of testis (thousands) between naïve and paired males of each species (prairie voles – Microtus ochrogaster, meadow voles – Microtus pennsylvanicus, and house mice – Mus musculus). Violin plots with individual data points represent data for each animal measured. Navy-colored bars represent the mean. Error bars indicate standard error. *p<0.05, **p<0.01, ***p<0.001, p-values between 0.05-0.1 are stated, “N.S.” indicates p>0.1.

## Results

First, summed testis weight, total mature sperm cell count, and mature sperm per mg of testis weight were compared between 22 naïve and 14 paired prairie voles, 15 naïve and 10 paired meadow voles, and 10 naïve and 10 paired house mice. Next, testicular histology was performed on a subset of each of the above groups, comparing 7 naïve and 8 paired prairie voles, 5 naïve and 5 paired meadow voles, and 3 naïve and 5 paired house mice.

### Summed Testis Weight: Multi-Species

Summed testis weights represent the combined weight of both testes per animal. Naïve prairie voles exhibited significantly smaller testes than paired prairie voles (0.463g ± 0.028g vs 0.651g ± 0.033g; *p*= 0.0052; t-statistic = -2.878; Figure 1A). There was no significant difference between naïve and paired meadow voles (0.765g ± 0.076g vs 0.657g ± 0.095g; *p*= 0.1669; t-statistic = 1.396; Figure 1A), nor between naïve house mice and paired house mice (0.187g ± 0.006g vs 0.199g ± 0.004g; *p*= 0.8933; t-statistic = -0.135; Figure 1A).

### Mature Sperm Cell Count: Multi-Species

Sperm counts presented are the summed number of sperm cells extracted from the cauda epididymides of both testes. Naïve prairie voles exhibited significantly lower mature sperm cell counts than paired prairie voles (93.2 ± 12.3 million vs 172 ± 22.3 million; *p*= 0.0043; t-statistic = -2.940; Figure 1B). There was no significant difference between naïve meadow voles or paired meadow voles (156 ± 28.3 million vs 121 ± 35.7 million; *p*= 0.2758; t-statistic = 1.098; Figure 1B), nor between naïve house mice and paired house mice (19.3 ± 1.73 million vs 20.8 ± 1.74 million; *p*= 0.9655; t-statistic = - 0.043; Figure 1B).

### Mature Sperm per mg of Testis: Multi-Species

Mature sperm per mg of testis represents the number of harvested mature sperm cells relative to the total testis mass per mg. Naïve prairie voles exhibited significantly less mature sperm cells per mg of testis weight compared to paired prairie voles (395 ± 34.7 thousand vs 471 ± 61.7 thousand; *p*= 0.0314; t-statistic = -2.192; Figure 1C). There was no significant difference between naïve and paired meadow voles (379 ± 51.7 thousand vs 334 ± 53.0 thousand; *p*= 0.5145; t-statistic = 0.655; Figure 1C), nor between naïve and paired house mice (204 ± 15.0 thousand vs 209 ± 15.9 thousand; *p*= 0.9445; t-statistic =-0.070; Figure 1C).

### Testicle Histology: Multi-Species

Tubule number, premature germ cells, Leydig cells, and Sertoli cells were quantified via histological examination. Tubule number and total number of premature germ cells can be seen in Figure 2. Figure 3 demonstrates the number of developing germ cells within each spermatogenic development stage (spermatogonia, spermatocytes, early spermatids, and late spermatids). Histology images represented of each experimental group can be seen in Figure 4. All histology data means, including *p*-values and t-statistics, is summarized in Table 1. Naïve prairie voles had significantly fewer tubules than paired prairie voles (Figure 2A). There was no significant difference in tubule number between naïve and paired meadow voles or house mice (Figure 2A). There was no significant difference in Leydig cell number between naïve and paired individuals of any species. There was no significant difference in Sertoli cell number between either naïve and paired prairie voles and house mice, however paired meadow voles had significantly more Sertoli cells than naïve meadow voles, despite displaying no increases in sperm production. There were no significant differences in the number of premature germ cells of any development stage between naïve and paired meadow voles or house mice (Figure 2B). Naïve prairie voles had significantly fewer spermatocytes, early spermatids, and late spermatids compared to paired prairie voles, but did not have significantly different numbers of spermatogonia (Figure 2B).

**Figure 2:**
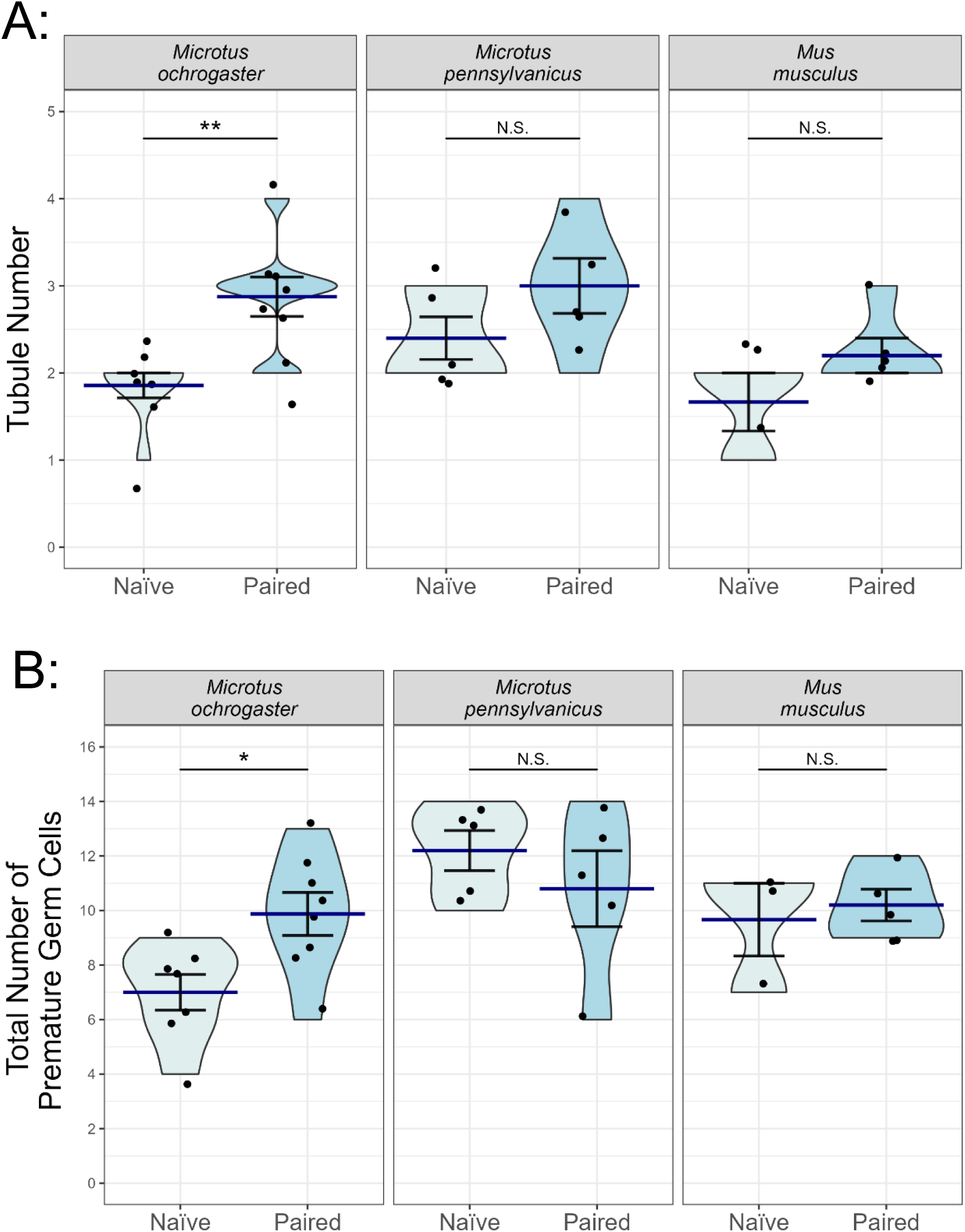
Comparison of A) the tubule number and B) total number of premature germ cells between naïve and paired males of each species (prairie voles – Microtus ochrogaster, meadow voles – Microtus pennsylvanicus, and house mice – Mus musculus). Violin plots with individual data points represent data for each animal measured. Navy-colored bars represent the mean. Error bars indicate standard error. *p<0.05, **p<0.01, ***p<0.001. p-values between 0.05-0.1 are stated, “N.S.” indicates p>0.1.

**Figure 3:**
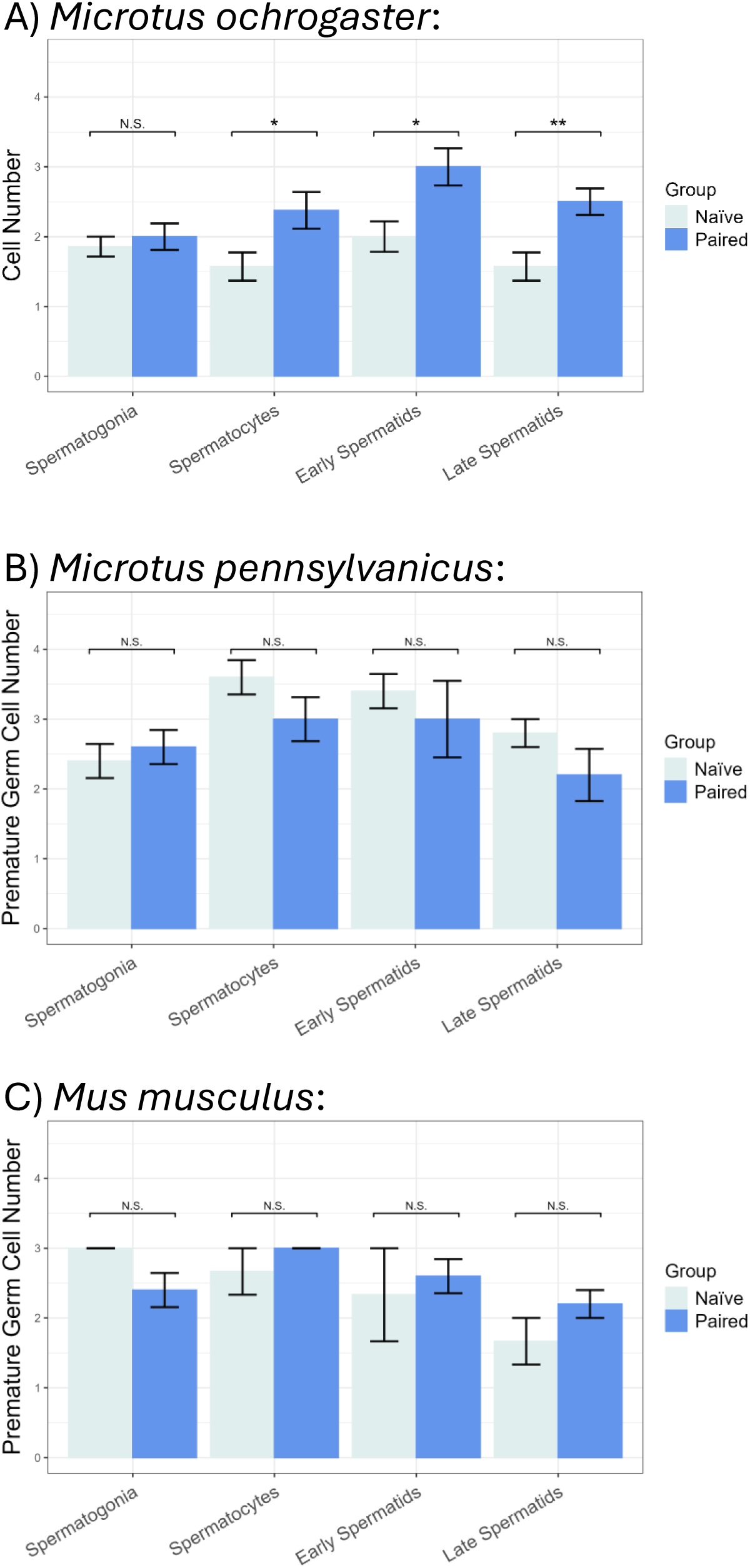
Comparison of cell number at each spermatogenic development stage between naïve and paired A) prairie voles (Microtus ochrogaster), B) meadow voles (Microtus pennsylvanicus), and C) lab mice (Mus musculus). Columns represent the mean of each group. Error bars indicate standard error. *p<0.05,**p<0.01,***p<0.001, N.S. indicates p>0.05.

**Figure 4:**
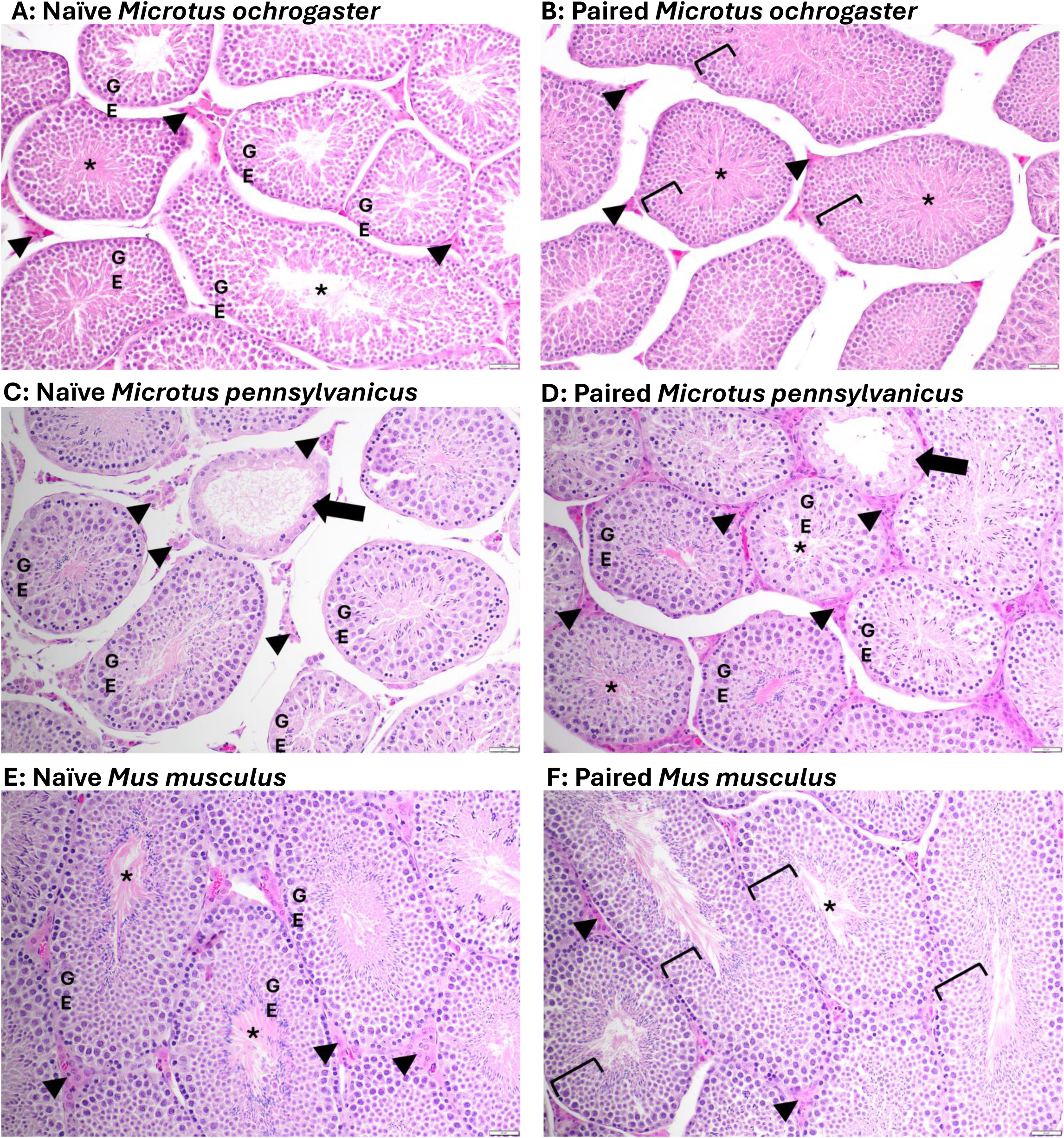
Testicle cross-sections from a A) naïve prairie vole (Microtus ochrogaster), B) paired prairie vole, C) naïve meadow vole (Microtus pennsylvanicus), D) paired meadow vole, E) naïve house mouse (Mus musculus), and F) paired house mouse. Lumen of seminiferous tubule (asterisk). Leydig cells in interstitial space (arrowheads). Germinal epithelium (left brackets or GE). Atrophic seminiferous tubule (arrows).

**Table 1:**
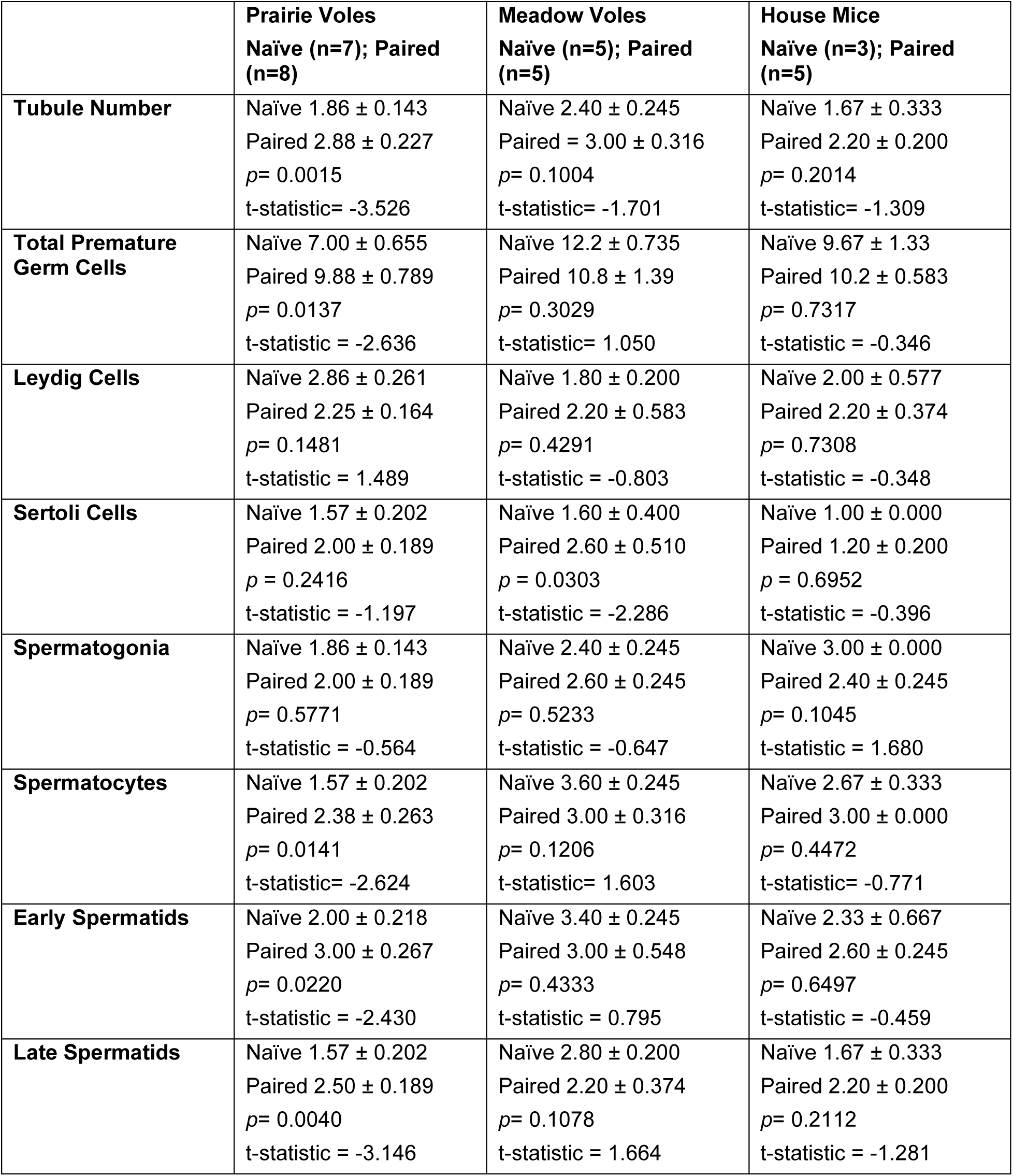
Comparison of testicle histology data means ± standard error between naïve and paired prairie voles, meadow voles, and house mice, including standard deviation, p-values, and t-statistics.

### Determining the Involvement of the Olfactory System in Novel Female-Induced Testicular Plasticity – Prairie Voles

The above data indicate that female pairing alone, in the absence of sperm competition, will induce an increase in testis weight and sperm production in prairie voles but not in meadow voles or house mice. To begin the investigation of possible mechanisms for this phenomenon, further analyses were performed on prairie voles with the addition of males regularly exposed only to soiled bedding from a novel female cage (the olfactory exposure group - OE), due to the importance of the olfactory system in both reproduction and pair-bond development (Decoster et al., 2023; Walum & Young, 2018). First, summed testis weight, total mature sperm cell count, and mature sperm per mg of testis weight were compared between 22 naïve, 14 paired, and 24 OE prairie voles. Next, testicular histology was performed on a subset of each group (7 naïve, 10 OE, and 8 paired). Finally, fecal testosterone metabolites were analyzed on 8 new individuals of each experimental group.

### Testis Mass and Sperm Count: Prairie Voles

When comparing the three prairie vole experimental groups, paired males exhibited significantly larger testes (0.651 ± 0.033g; *p*= 0.0005) than naïve males (0.463 ± 0.028g; Figure 5A). OE males averaged between the two groups (0.564 ± 0.030g; Figure 5A), with testes statistically larger than naïve males (*p*= 0.0404) but not statistically different from the paired males (*p*= 0.1522). Paired males exhibited significantly higher mature sperm counts (172 ± 22.3 million; *p*= 0.0008; t-statistic = - 3.861) than naïve males (93 ± 12.3 million; Figure 5B). OE males (143 ± 8.86 million) averaged between naïve and paired males but were not statistically different from either group (naïve-OE *p*= 0.0819, t-statistic = -2.188; OE-paired *p*= 0.1203, t-statistic = - 2.005; Figure 5B). Paired males trended (*p*= 0.0802) toward more mature sperm cells per mg of testis weight (522 ± 61.7 thousand) compared to naïve males (395 ± 34.7 thousand; Figure 5C), however this was not statistically significant. OE males (471 ± 26.3 thousand) averaged between naïve and paired males but did not statistically differ from either group (naïve-OE *p*= 0.2812, t-statistic = -1.539; OE-paired *p*= 0.6521, t-statistic = -0.885; Figure 5C).

**Figure 5:**
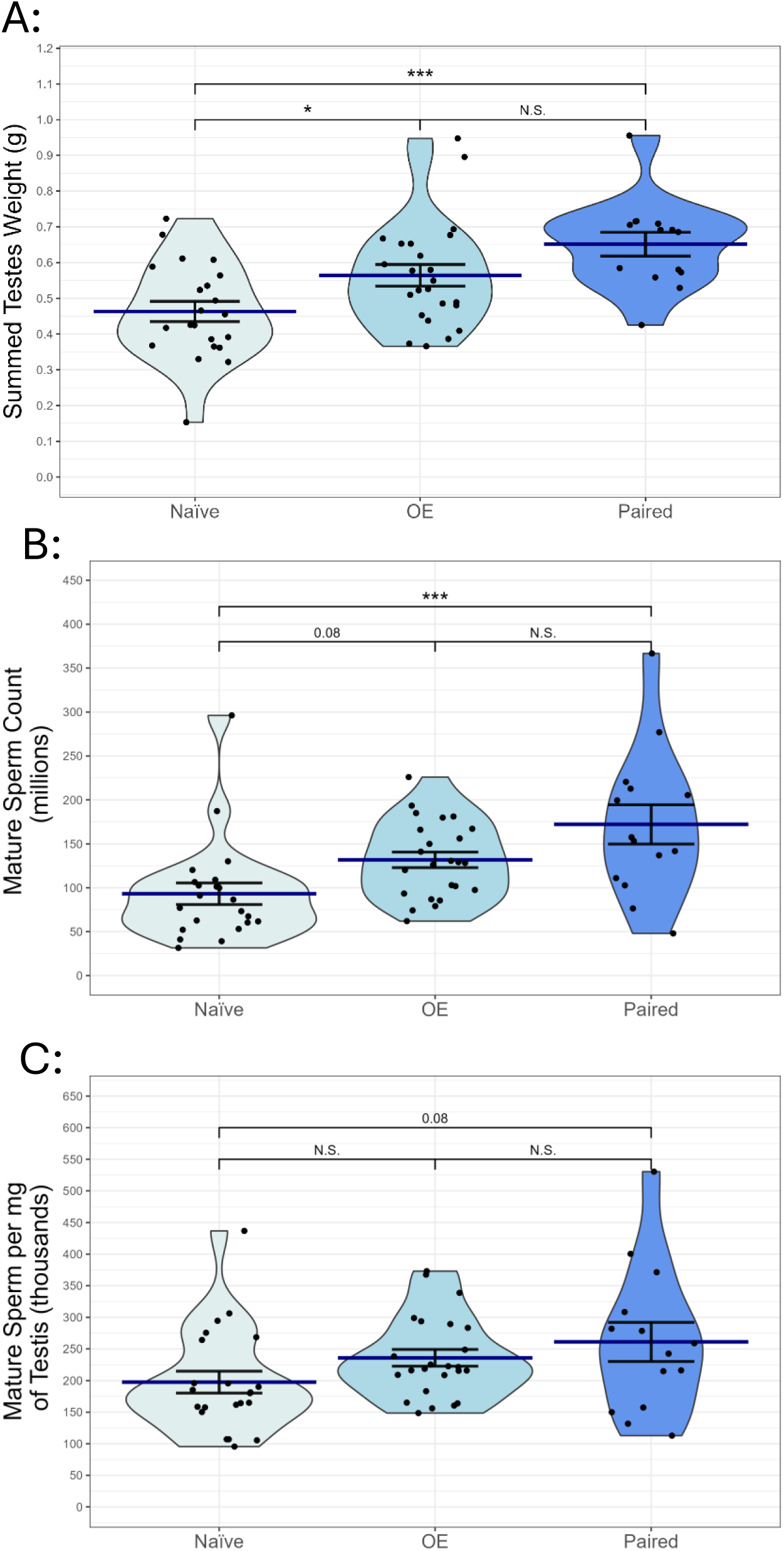
Comparison of A) summed testis weight (g), B) mature sperm count (millions), and C) mature sperm per mg of testis (thousands) between naïve, OE, and paired prairie vole (Microtus ochrogaster) males. Violin plots with individual data points represent the full data distribution. Navy-colored bars represent the mean. Error bars indicate standard error.

### Testicle Histology: Prairie Voles

As was done for the multi-species comparison, tubule number, premature germ cells, Leydig cells, and Sertoli cells were compared between the three prairie vole groups. All histology data – including statistical analysis – is summarized in Table 2. There were no significant differences in the number of Leydig or Sertoli cells between any group. Naïve males had significantly fewer tubules than OE and paired males, while OE and paired males had statistically similar tubule numbers (Figure 6A). Naïve males had significantly fewer premature germ cells than paired males. OE males trended to a similar number of premature germ cells as paired males but were not significantly different from either naïve or paired males (Figure 6B. When the premature germ cells are categorized into each spermatogenic development stage (spermatogonia, spermatocyte, early spermatids, and late spermatids), all groups began with a similar number of spermatogonia. OE and paired males exhibit greater numbers of spermatocytes, early spermatids, and late spermatids than naïve males (Figure 7). Histology images representive of each experimental group can be seen in Figure 8.

**Figure 6:**
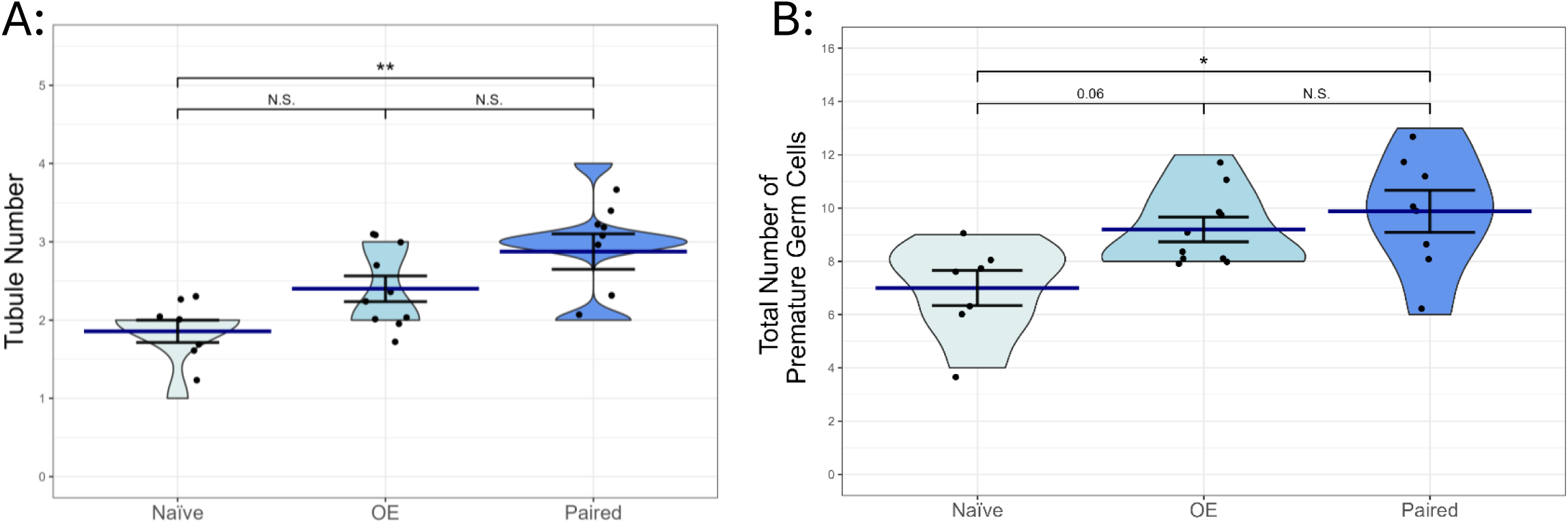
Comparison of A) the tubule number and B) total number of premature germ cells between naïve, OE, and paired prairie vole (Microtus ochrogaster) males. Violin plots with individual data points represent data for each animal measured. Navy-colored bars represent the mean. Error bars indicate standard error. *p<0.05, **p<0.01, ***p<0.001, p-values between 0.05-0.1 are stated, “N.S.” indicates p>0.1.

**Figure 7:**
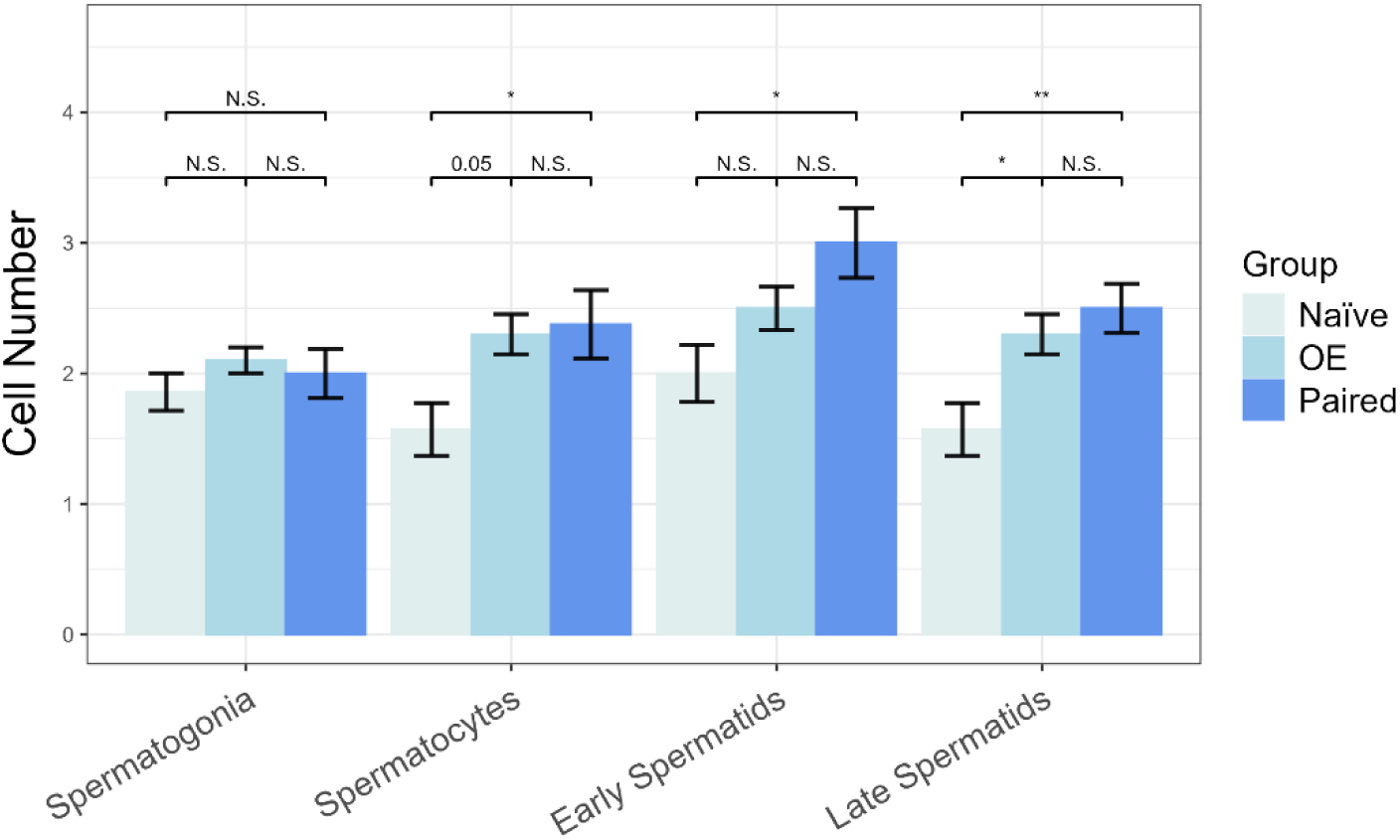
Comparison of cell number at each spermatogenic development stage between naïve, OE, and paired prairie vole (Microtus ochrogaster) males. Columns represent the mean of each group. Error bars indicate standard error. *p<0.05, **p<0.01, ***p<0.001, p-values between 0.05-0.1 are stated, “N.S.” indicates p>0.1.

**Figure 8:**
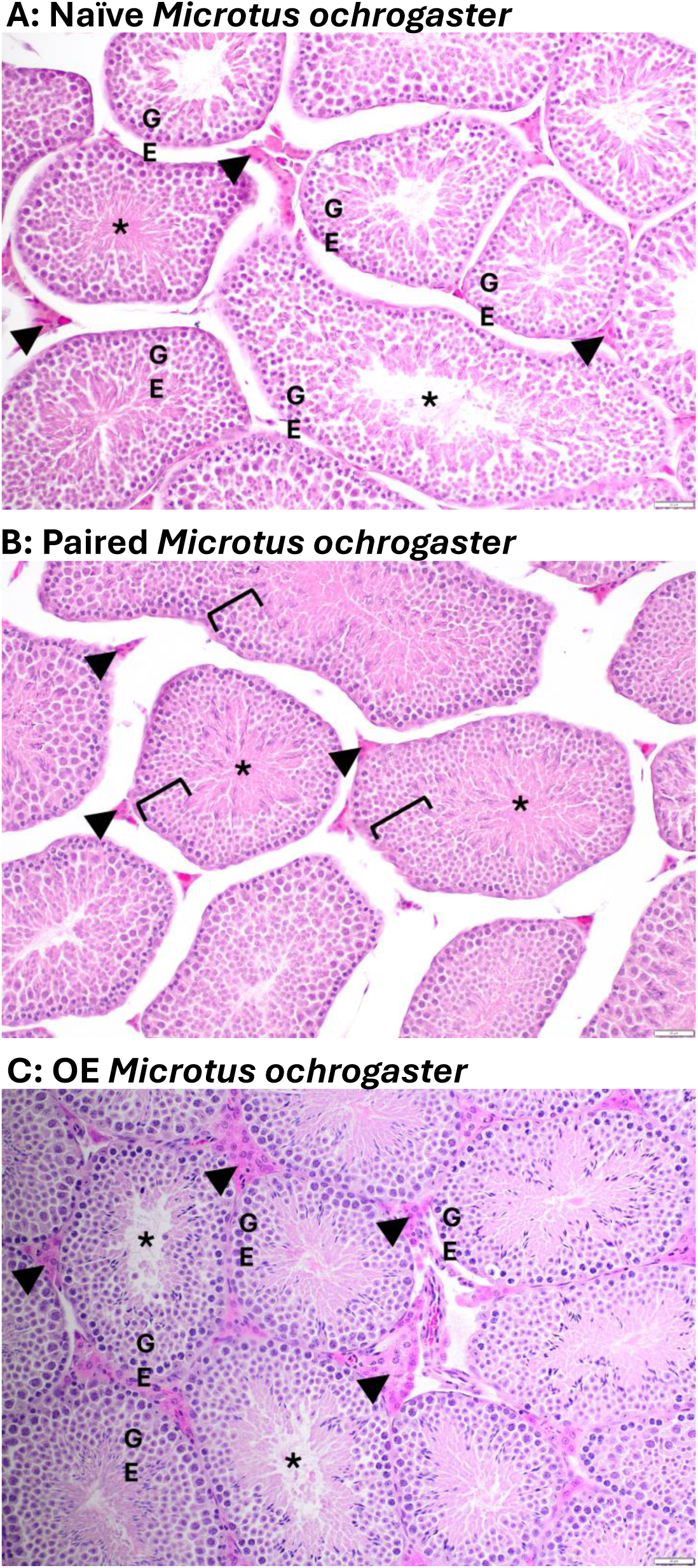
Testicle cross-sections from a A) naïve prairie vole (Microtus ochrogaster), B) paired prairie vole, and C) OE prairie vole. Lumen of seminiferous tubule (asterisk). Leydig cells in interstitial space (arrowheads). Germinal epithelium (left brackets or GE).

**Table 2:** Comparison of testicular histology data means ± standard error between naïve, OE, and paired prairie voles, including p-values and t-statistics.

|  | **Naïve**<br>**(n = 7)** | **OE**<br>**(n = 10)** | **Paired**<br>**(n = 8)** | **t-statistic** | **p-values** |
| --- | --- | --- | --- | --- | --- |
| **Tubule Number** | 1.86 ± 0.143 | 2.40 ± 0.163 | 2.88 ± 0.227 | Naïve-Paired: -3.725<br>Naïve-OE: -2.086<br>OE-Paired: -1.897 | Naïve-Paired: 0.0032<br>Naïve-OE: 0.1158<br>OE-Paired: 0.1633 |
| **Total Premature Germ Cells** | 7.00 ± 0.655 | 9.20 ± 0.467 | 9.88 ± 0.789 | Naïve-Paired: -3.061<br>Naïve-OE: -2.460<br>OE-Paired: -0.784 | Naïve-Paired: 0.0152<br>Naïve-OE: 0.0557<br>OE-Paired: 0.7165 |
| **Leydig Cells** | 2.86 ± 0.261 | 2.30 ± 0.213 | 2.25 ± 0.164 | Naïve-Paired: 1.892<br>Naïve-OE: 1.823<br>OE-Paired: 0.170 | Naïve-Paired: 0.1646<br>Naïve-OE: 0.1853<br>OE-Paired: 0.9842 |
| **Sertoli Cells** | 1.57 ± 0.101 | 2.00 ± 0.149 | 2.00 ± 0.189 | Naïve-Paired: -1.625<br>Naïve-OE: -1.706<br>OE-Paired: 0.00 | Naïve-Paired: 0.2566<br>Naïve-OE: 0.2252<br>OE-Paired: 1.00 |
| **Spermatogonia** | 1.86 ± 0.143 | 2.10 ± 0.100 | 2.00 ± 0.189 | Naïve-Paired: -0.668<br>Naïve-OE: -1.192<br>OE-Paired: 0.510 | Naïve-Paired: 0.7843<br>Naïve-OE: 0.4699<br>OE-Paired: 0.8672 |
| **Spermatocytes** | 1.57 ± 0.202 | 2.30 ± 0.153 | 2.38 ± 0.263 | Naïve-Paired: -2.626<br>Naïve-OE: -2.501<br>OE-Paired: -0.267 | Naïve-Paired: 0.0394<br>Naïve-OE: 0.0512<br>OE-Paired: 0.9614 |
| **Early Spermatids** | 2.00 ± 0.218 | 2.50 ± 0.167 | 3.00 ± 0.267 | Naïve-Paired: -3.108<br>Naïve-OE: -1.632<br>OE-Paired: -1.696 | Naïve-Paired: 0.0136<br>Naïve-OE: 0.2536<br>OE-Paired: 0.2291 |
| **Late Spermatids** | 1.57 ± 0.202 | 2.30 ± 0.153 | 2.50 ± 0.189 | Naïve-Paired: -3.490<br>Naïve-OE: -2.876<br>OE-Paired: -0.820 | Naïve-Paired: 0.0056<br>Naïve-OE: 0.0229<br>OE-Paired: 0.6947 |

### Fecal Testosterone: Prairie Voles

There were no statistical differences between prairie vole experimental groups in fecal testosterone metabolite concentration (naïve-paired *p*= 0.3711, t-statistic = 1.376; naïve-OE *p*= 0.6712, t-statistic = 0.859; OE-paired *p*= 0.8642, t-statistic = 0.517; Figure 9). Means and standard error were as follows: naïve (5.45 ± 0.150ng/g), OE (5.06 ± 0.314ng/g), and paired (4.84 ± 0.418ng/g).

**Figure 9:**
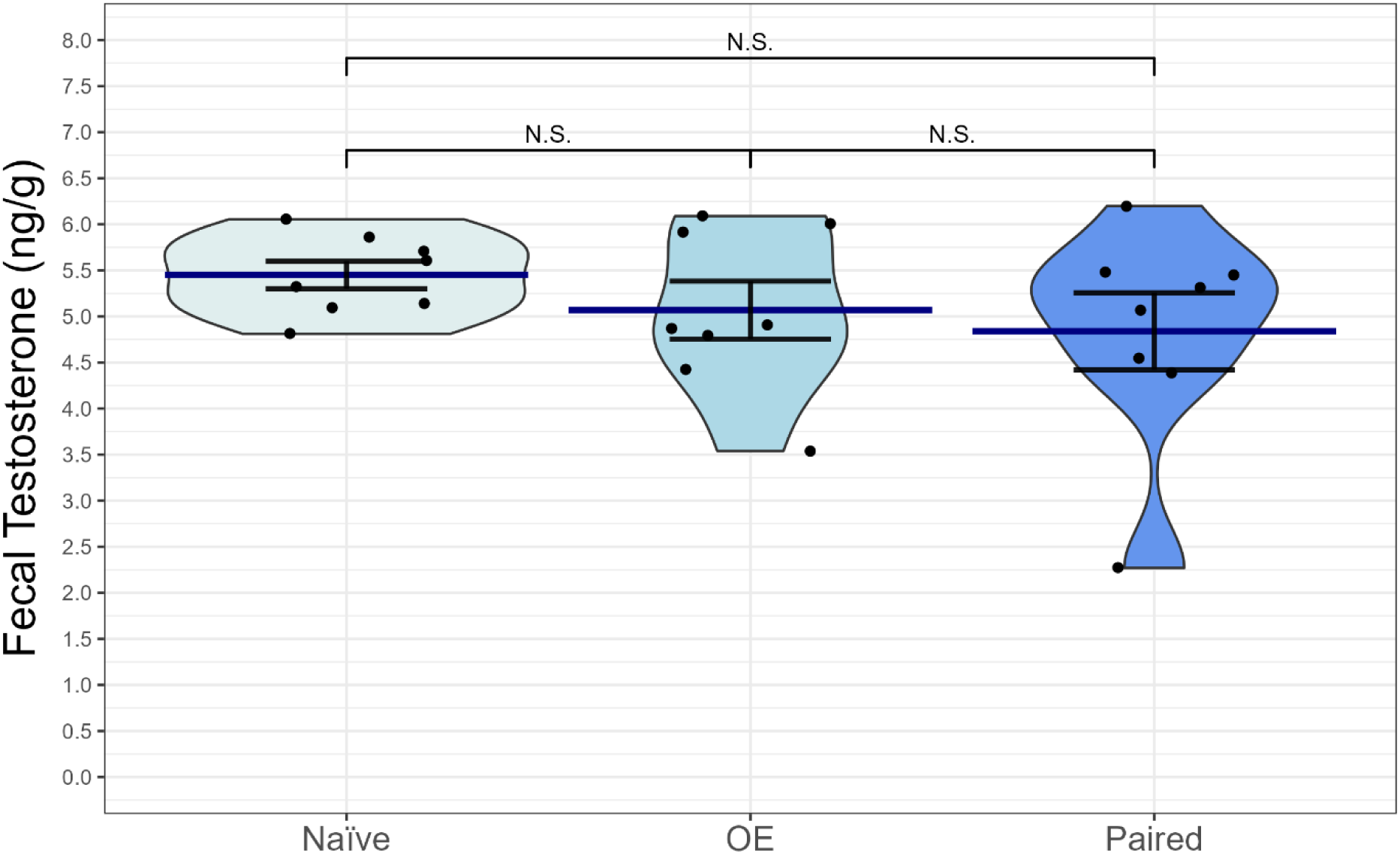
Comparison of fecal testosterone metabolites between naïve, OE, and paired prairie vole (Microtus ochrogaster) males. Violin plots with individual data points represent data for each animal measured. Navy-colored bars represent the mean. Error bars indicate standard error. *p<0.05, **p<0.01, ***p<0.001, p-values between 0.05-0.1 are stated, “N.S.” indicates p>0.1.

### Comparing Prairie Voles to Meadow Voles: Sperm Competition Theory

Sperm competition theory states that male individuals of species with lower rates of female promiscuity exhibit smaller testes and less sperm production than the males of closely related species with higher rates of female promiscuity (Lüpold et al., 2020). To examine the applicability of this theory in prairie voles and meadow voles, comparisons between summed testis weight and mature sperm count were made between 22 naïve prairie voles and 15 naïve meadow voles and 14 paired prairie voles and 10 paired meadow voles.

Naïve meadow voles exhibited significantly (*p*= 0.0001; t-statistic = -4.193) larger testes (0.765 ± 0.076g) than naïve prairie voles (0.463 ± 0.028g; Figure 8A). Naïve meadow voles also exhibited significantly (p= 0.0375; t-statistic = -2.129) more mature sperm (156 ± 28.3 million) than naïve prairie voles (93 ± 12.3 million; Figure 8B). However, there were no significant differences between paired prairie vole males and paired meadow vole males in either testis weight (0.651 ± 0.033g vs 0.657 ± 0.095g; p= 0.9511; t-statistic = -0.062; Figure 10A) or sperm production (172 ± 22.3 million vs 121 ± 35.7 million; p= 0.1768; t-statistic = 1.367; Figure 10B).

**Figure 10:**
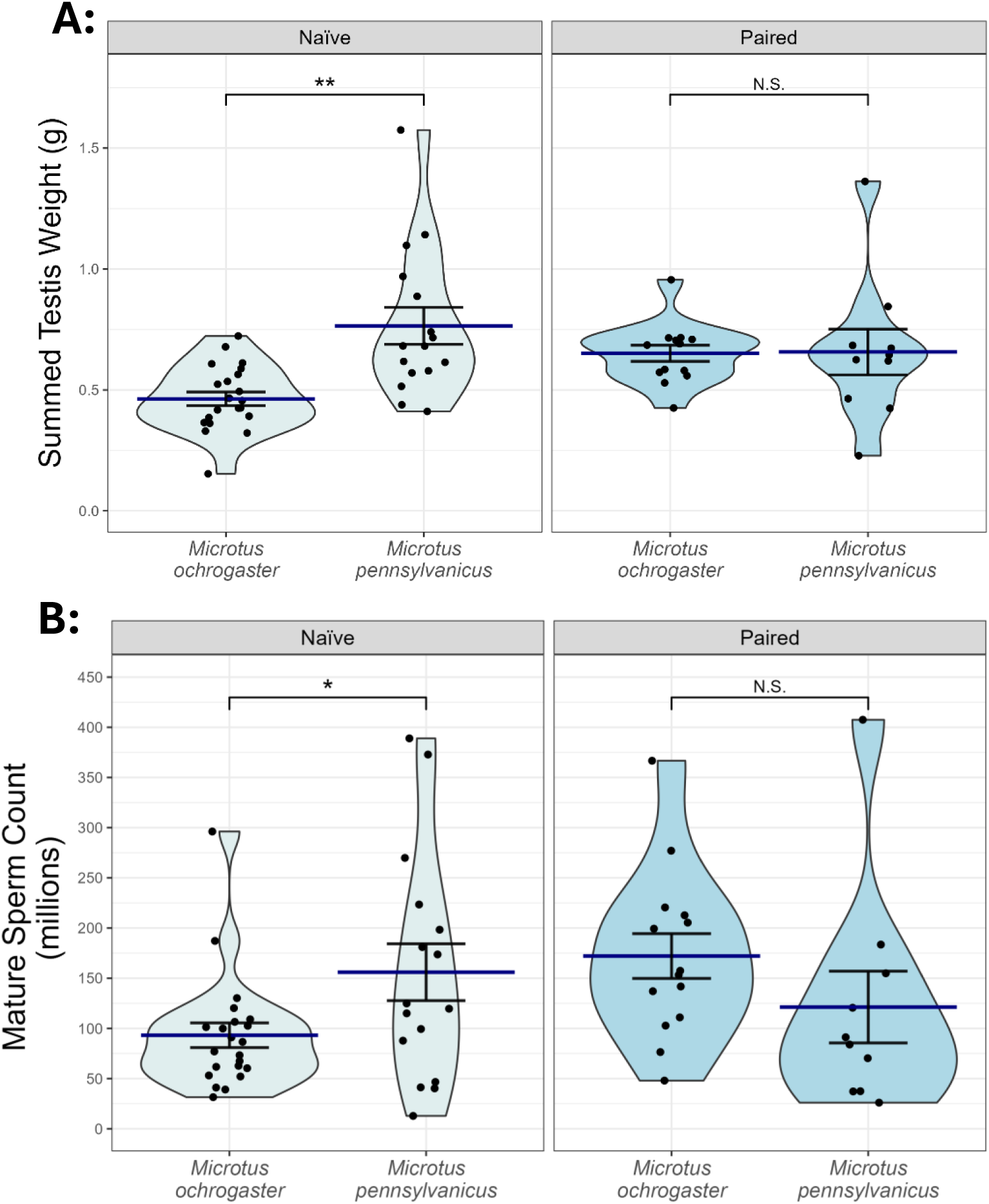
Comparison of A) summed testis weight (g) and B) mature sperm count (millions) between naïve prairie voles (Microtus ochrogaster) and meadow voles (Microtus pennsylvanicus) and paired prairie voles and meadow voles. Violin plots with individual data points represent data for each animal measured. Navy-colored bars represent the mean. Error bars indicate standard error. *p<0.05, **p<0.01, ***p<0.001, p-values between 0.05-0.1 are stated, “N.S.” indicates p>0.1.

### Examining for Weight and Age Effects for all Groups and Species

Body weight had no significant impact on summed testis weight or mature sperm count in naïve prairie voles (*p*= 0.473, R^2^= -0.0335; *p*=0.7045, R^2^= -0.0461), OE prairie voles (*p*= 0.6169, R^2^= -0.0334; *p*=0.5597, R^2^= -0.0291), paired prairie voles (*p*= 0.5658, R^2^= - 0.0572; *p*=0.856, R^2^= -0.0875), naïve meadow voles (*p*= 0.0770, R^2^= 0.4797; *p*= 0.2395, R^2^= 0.1535), or paired meadow voles (*p*= 0.6532, R^2^= -0.2319; *p*= 0.5685, R^2^= - 0.1737). Body weight was positively correlated with summed testis weight, but not mature sperm count in naïve mice (*p*= 0.0002, R^2^= 0.8090; *p*= 0.0801, R^2^= 0.2509). Body weight was not positively correlated with summed testis weight but was positively correlated with mature sperm count in paired mice (*p*= 0.4116, R^2^= -0.0307; *p*= 0.0130, R^2^= 0.5543).

Age had no significant impact on summed testis weight or mature sperm count in naïve prairie voles (*p*= 0.8359, R^2^= -0.0477; *p*=0.6146, R^2^= -0.0365), OE prairie voles (*p*= 0.9494, R^2^= -0.0453; *p*=0.6875, R^2^= -0.0376), naïve meadow voles (*p*= 0.8901, R^2^= - 0.0699; *p*= 0.9205, R^2^= -0.0706), paired meadow voles (*p*= 0.8493, R^2^= -0.1196; *p*= 0.3286, R^2^= 0.0091), or paired mice (*p*= 0.2117, R^2^= 0.0856; *p*= 0.6036, R^2^= -0.0854). In paired prairie voles, age had no impact on summed testis weight but exhibited correlation with mature sperm count (p=0.706, R^2^= -0.07003; p=0.0143, R^2^= 0.3564). However, this was driven by one older individual with a high sperm count. Age was positively correlated with both summed testis weight and mature sperm count in naïve mice (*p*=0.0046, R^2^= 0.6112; *p*= 0.0505, R^2^= 0.3228).

Overall, age and weight should not have an impact between group and species comparisons. Naïve mice were the only group to exhibit a consistent effect of age and weight; however, the individuals of this group were already quite similar in age and weight.

### Effects of Female Exposure Length for Female-Exposed Experimental Groups – All Species

All data was gathered post female exposure after a minimum of one spermatogenic cycle. However, the length of female exposure varied greatly, from only one spermatogenic cycle for some individuals to many months for others. Therefore, potential impacts of pairing length on summed testis weight and mature sperm count were examined. Length of female exposure produced no significant effect on either summed testis weight nor mature sperm count in OE prairie voles (*p*= 0.1202, R^2^= 0655; *p*= 0.2728, R^2^= 0.0114), paired prairie voles (*p*= 0.8532, R^2^= -0.0801), paired meadow voles (*p*= 0.2299, R^2^= 0.0711; *p*= 0.5007, R^2^= -0.0592), or paired house mice (*p*= 0.1258, R^2^= 0.1759; *p*= 0.6044, R^2^= -0.0855).

## Discussion

Male prairie voles paired monogamously with a female exhibit 41% larger testes and 85% more mature sperm cells than naïve males. These changes are accompanied by numerous differences in testicular histology. This pattern is not observed in meadow voles or house mice, which exhibit no significant differences between naïve or paired individuals (except for Sertoli cells in meadow voles). In general, perceived sperm competition can increase sperm production and alter sperm investment amongst numerous male mammals (Burger et al., 2015; delBarco-Trillo & Ferkin, 2006; Dixson & Anderson, 2004; Firman et al., 2018; Ramm & Stockley, 2009b). However, research on the effects of female exposure alone on gonad size or function across different mammal species is sparse, which makes this finding in prairie voles intriguing. The OE prairie vole group revealed that this plasticity in prairie voles is at least partially modulated by the olfactory system. In some rodent studies using promiscuous species, exposure to only olfactory cues from male rivals triggered changes in male reproductive physiology (delBarco-Trillo & Ferkin, 2006; Ramm & Stockley, 2009b). Perhaps this phenomenon in prairie voles is an evolutionary adaptation of this existing circuitry in response to the development of their social behavior. Additionally, the entirety of this female-induced change occurs within one cycle of spermatogenesis (∼29 days). Longer lengths of female exposure do not further increase testis size nor mature sperm counts.

All experimental prairie vole groups exhibit similar numbers of spermatogonia, but differ in spermatocytes, early spermatids, and late spermatids, with female-exposed males displaying higher numbers. Spermatogenesis is regulated in the testes via complex molecular interactions largely modulated by luteinizing hormone (LH) and follicle stimulating hormone (FSH) and their effects on Leydig and Sertoli cells, respectively (Oduwole et al., 2018; Smith & Walker, 2014). LH and FSH are regulated by a system of gonadotropin releasing hormone (GnRH) neurons which project to the median eminence, the majority of which reside in the medial preoptic area of the hypothalamus (Casteel & Singh, 2024; Kaprara & Huhtaniemi, 2018). The kisspeptin network also has significant involvement in regulating GnRH communication (Sobrino et al., 2022). While numerous studies exhibit strong effects of social experiences/environments on the GnRH system, the specific pathways and neurotransmitters regulating these phenomena are largely uncharacterized (Maruska & Fernald, 2011b).

We did not find statistical differences in fecal testosterone metabolites between naïve, OE, and paired males. Wang et al. (1993) reported that three days of male-female cohabitation raised plasma testosterone in male prairie voles, but not in male meadow voles (Wang et al., 1994). The lack of differences in fecal testosterone metabolites in the present study may be due to differences in free or bound hormone quantification in fecal vs plasma measurements, or perhaps a greater length of female or female odor exposure time in our female-exposed males. The observed changes in spermatogenesis between prairie vole experimental groups suggest a likelihood of differences in testicular testosterone, but this was not measured in the present study. Testosterone plays a key role in the regulation of spermatogenesis and rodent testicular tissue contains much higher levels of testosterone than is found in serum (Smith & Walker, 2014). When Leydig cells produce testosterone, Sertoli cells secrete androgen binding protein (ABP) into the seminiferous tubules in response (Hammond & Hogeveen, 2003). ABP is thought to increase testosterone concentrations in the seminiferous tubules and epididymal cells to facilitate spermatogenesis, however the location, relative expression, and role of ABP appears to vary somewhat between mammalian species (Hammond & Hogeveen, 2003; Robaire & Hinton, 2015). It would be ideal to examine testosterone and ABP concentrations and expressions in the seminiferous tubules and epididymal tissue of naïve and female-exposed prairie voles to better understand the involvement of androgens in their testes.

The OE prairie vole males most often statistically resembled paired prairie voles, but consistently averaged between the naïve and paired groups in nearly all measured variables. A larger sample size of paired males may more definitively determine if there are gonadal differences between OE and paired prairie vole males. Ophir and delBarcillo-Trillo (2007) found that, in a semi-natural field enclosure, prairie vole males that pair bonded with a female (AKA “resident” males) exhibited larger testes and more sperm than males which did not pair bond, but still experienced regular females encounters and copulatory opportunities (AKA “wandering” males) (Ophir & delBarco-Trillo, 2007). This was interpreted to be a function of female mate preference for a larger anogenital distance (AGD). Notably, the male-female pairs in the present study were randomly mated, eliminating female mate choice. In this case, the paired males still exhibited significantly larger testes than naïve males, but not significantly larger than our OE males. It is therefore possible that perceived sperm competition in addition to pair bonding may further increase testis size and sperm production in paired prairie vole males, or that pair bonding could induce a greater increase in testis mass and sperm production if a larger sample size is examined. This would provide paired prairie vole males an additional advantage against potential cuckoldry and potentially increase their own attractiveness to their bonded mate via an enlarged AGD. Further work should be done to examine additional effects of perceived sperm competition in paired prairie vole males.

It is possible that the evolutionary history leading to modern prairie vole social structure has resulted in alterations to male reproductive physiology which provide reproductive advantages that suit social (but often not genetic) monogamy alongside family and colony living. Many prairie voles remain in their natal burrows for their entire lives and both sexes are known to act as alloparents (Finton et al., 2022; McGuire et al., 1993). In this social environment, opportunities to reproduce may be less plentiful than those beyond the natal nest (McGuire et al., 1993). Female prairie voles remain in an anestrus state until they encounter unfamiliar conspecific males. They will not begin to cycle solely in the presence of their fathers or familiar male siblings (C. Carter, 1987; C. S. Carter et al., 1980). Thus, our observation that naïve males exhibit smaller testes and less sperm than OE and paired males may represent the male counterpart of this female reproductive suppression. If or when these young males begin venturing father from their home burrows, olfactory, social, and reproductive encounters with novel conspecifics may trigger an increase in testis size and sperm production.

The strength of novel female exposure on male prairie vole reproductive physiology is further reflected by comparing the testis size and sperm count of prairie voles and meadow voles. Sperm competition theory proposes that male individuals of species with lower rates of female promiscuity exhibit smaller testes and less sperm production than the males of closely related species with higher rates of female promiscuity, which has been supported by several studies (Csanády et al., 2019; Lüpold et al., 2020; Simmons et al., 2004). Although this theory is consistent with observations between naïve prairie voles and meadow voles, there is no difference in testis size or sperm production between paired prairie voles and meadow voles. This is a testament to both the strength of plasticity induced by female exposure in prairie voles and the complexity of promiscuity that exists within socially monogamous species.

Prairie vole colonies across North America exhibit some variability in social behavior (Ortiz et al., 2022). The data acquired in this study should ideally be replicated via both lab and field data for prairie vole colonies in different locations to determine if this phenomenon is consistent geographically. Since it appears that testicular plasticity in prairie voles is strongly modulated by the social brain, future work should investigate the brain regions and neurotransmitters involved in regulating this plasticity. Additionally, this study sought to begin examining prairie vole testicular plasticity within a monogamy-promiscuity framework with the inclusion of meadow voles and house mice. To determine if this relationship applies broadly to other mammalian taxa, more comparisons must be made between closely related species with differences in mating and parental strategy. Although there is an established relationship between female promiscuity and male testis size between species (Kramer & Russell, 2015; Schacht & Kramer, 2019; van der Horst & Maree, 2014), the ways in which testicular plasticity is modulated within socially monogamous species remain largely unexplored.

## Conclusion

Prairie voles exhibit testicular plasticity and adaptive sperm production in response solely to both direct and indirect female exposure – a phenomenon not found in meadow voles or house mice. Statistical similarities between OE and paired prairie vole males suggest this plasticity is modulated via undetermined social and olfactory neural pathways communicating with the GnRH neuronal network. It is possible this manifestation of testicular plasticity in prairie voles arose as an evolutionary adaptation to social monogamy and paternal care to improve male reproductive success depending on individual mating strategy. Whether or not similar a phenomenon occurs in other socially monogamous, biparental species warrants further investigation.

## Credit Authorship Contribution Statement

**Jessica Hurd**: Original draft preparation, conceptualization, visualization, investigation, methodology, validation, formal analysis, data curation. **Yurika Watanabe, Gracie Toben,** and **Casey Sergott**: Data curation, reviewing and editing. **Alexandra Ford** and **Craig Miller**: Data curation and tissue evaluation (histology), writing. **Dale Kelley**: Conceptualization, resource provision, methodology, validation. **Zoe Donaldson**: Resource provision, funding acquisition. **Elizabeth McCullagh**: Resource provision, conceptualization, project administration, supervision. All authors contributed to manuscript reviewing and editing.

## Funding and Acknowledgements

This work was supported by the National Institute of Health [NIH R01 MH125423 (ZRD), NIH DP2 MH119427 +S1 (ZRD), NIH 5T32 GM140953 (JAH)], NIH NIGMS P30GM149368] and National Science Foundation [NSF CAREER IOS-2045348 (ZRD) and NSF IOS-1827790 (ZRD)]. The content is solely the responsibility of the authors and does not necessarily represent the official views of the National Institutes of Health or National Science Foundation

We thank Shay Nguyen for work on prairie vole data collection, Dr. Jennifer Grindstaff for providing equipment for the hormone assays, Dr. Phoebe Edwards for support with calibrating and troubleshooting the hormone assays, and Dr. Dani Kirsch for support with statistical analysis in R Studio.

## Data Availability Statement

Data will be made available on request.

